# Solar simulated ultraviolet radiation inactivates HCoV-NL63 and SARS-CoV-2 coronaviruses at environmentally relevant doses

**DOI:** 10.1101/2021.06.25.449831

**Authors:** Georg T. Wondrak, Jana Jandova, Spencer J. Williams, Dominik Schenten

**Affiliations:** Department of Pharmacology and Toxicology, College of Pharmacy and UA Cancer Center, University of Arizona, Tucson, Arizona; Department of Immunobiology, College of Medicine, University of Arizona, Tucson, Arizona

**Keywords:** Coronavirus, HCoV-NL63, SARS-CoV-2, Solar simulated ultraviolet radiation, Viral inactivation

## Abstract

The germicidal properties of short wavelength ultraviolet C (UVC) light are well established and used to inactivate many viruses and other microbes. However, much less is known about germicidal effects of terrestrial solar UV light, confined exclusively to wavelengths in the UVA and UVB regions. Here, we have explored the sensitivity of the human coronaviruses HCoV-NL63 and SARS-CoV-2 to solar-simulated full spectrum ultraviolet light (sUV) delivered at environmentally relevant doses. First, HCoV-NL63 coronavirus inactivation by sUV-exposure was confirmed employing (*i*) viral plaque assays, (*ii*) RT-qPCR detection of viral genome replication, and (*iii*) infection-induced stress response gene expression array analysis. Next, a detailed dose-response relationship of SARS-CoV-2 coronavirus inactivation by sUV was elucidated, suggesting a half maximal suppression of viral infectivity at low sUV doses. Likewise, extended sUV exposure of SARS-CoV-2 blocked cellular infection as revealed by plaque assay and stress response gene expression array analysis. Moreover, comparative (HCoV-NL63 versus SARS-CoV-2) single gene expression analysis by RT-qPCR confirmed that sUV exposure blocks coronavirus-induced redox, inflammatory, and proteotoxic stress responses. Based on our findings, we estimate that solar ground level full spectrum UV light impairs coronavirus infectivity at environmentally relevant doses. Given the urgency and global scale of the unfolding SARS-CoV-2 pandemic, these prototype data suggest feasibility of solar UV-induced viral inactivation, an observation deserving further molecular exploration in more relevant exposure models.

## 1. Introduction

The germicidal properties of short wavelength ultraviolet C (UVC) light are well established and widely used to inactivate many viruses and other microbes, and virucidal activity of solar UVC targeting pathogenic coronaviruses has been explored in much detail before [1–3]. Given the urgency and global scale of the unfolding SARS-CoV-2-caused COVID-19 pandemic, UV-induced inactivation of coronaviruses including SARS-CoV-2 has reemerged as a matter of much contemporary research interest [2–8]. Indeed, recently, rapid and complete inactivation of SARS-CoV-2 by UVC has been substantiated experimentally, and virucidal UVC light sources (254 nm emission) are used for surface disinfection and decontamination [5,8]. Moreover, far UVC (222 nm) has attracted considerable attention due to its potent virucidal activity [2]. However, much less is known about germicidal (and coronavirus-directed) effects of terrestrial (ground level) solar UV light, a matter of much interest given the airborne spread of coronaviruses including SARS-CoV-2 [2,6]. UVC (**<** 290 nm) is not present in the solar spectrum reaching the Earth’s surface, and most of solar UV energy incident on the skin is from the UVA region (**>**95%; from 320–400 nm). Remarkably, the UVB (290–320 nm) proportion of total solar UV-flux received by skin can be well below 2% depending on the solar angle, which determines the atmospheric light path length and thereby the degree of ozone-filtering and preferential Rayleigh scattering of short wavelength UV light [9].

Recently, the role of ground level (environmentally relevant) solar UV has been explored in the context of SARS-CoV-2 disinfection, and a role of solar UVB in human coronavirus inactivation has been substantiated based on atmospheric and geophysical simulations [2,6,10,11]. Specifically, inactivation times of SARS coronaviruses exposed to environmental photons with wavelengths between 290-315 nm have been calculated using OMI (ozone monitoring instrument) satellite data for the sunlit earth [10]. Moreover, recent research has demonstrated that simulated sunlight rapidly inactivates SARS-CoV-2 on surfaces including human saliva when exposed to simulated sunlight representative of the summer solstice at 40 °N latitude at sea level on a clear day [10]. Also, indirect effects of solar UVB exposure in reducing COVID-19 deaths have been substantiated, potentially mediated by UVB-driven cutaneous vitamin D synthesis, among other factors [12–14]. In addition, a role of solar UVA photons in the inactivation of coronaviruses has been proposed [7].

Given the complexity of virucidal activity as a function of spectral composition from ultraviolet to infrared, a topic recently reviewed by various authors, a more detailed knowledge and direct evidence of solar UV-induced coronavirus inactivation (achievable at ground level and environmentally relevant doses) would offer improved options that inform decisions at the basic research, clinical care, and public health levels [2,6,8]. Here, for the first time, we have explored the sensitivity of the human coronaviruses HCoV-NL63 and SARS-CoV-2 to solar simulated ultraviolet light (sUV). Our findings suggest that solar UV delivered at environmentally relevant dose levels inactivates HCoV-NL63 and SARS-CoV-2 coronaviruses with pronounced blockade of infectivity protecting mammalian host cells.

## 2. Materials and Methods

### 2.1. Chemicals

All chemicals were purchased from Sigma Aldrich (St. Louis, MO, USA).

### 2.2. Mammalian cell culture, viral propagation, and target cell infection

As established viral target cells infected by HCoV-NL63 and SARS-CoV-2, Calu-3 human metastatic lung epithelial adenocarcinoma (HTB-55), Caco-2 human colorectal epithelial adenocarcinoma (HTB-37) and Vero normal epithelial monkey kidney (CCL-81) cells (all from ATCC, Manassas, VA, USA) were maintained according to published standard procedures [15–18]. In brief, all cells (Calu-3, Caco-2 and Vero) were cultured in Eagle’s Minimum Essential Medium (MEM) medium (Corning, Manassas, VA) supplemented with 10% bovine calf serum (BCS, HyClone™ Laboratories, Logan, UT). Coronavirus HCoV-NL63 (NR-470) and its genomic RNA (NR-44105) were obtained from BEI Resources (NIAID, NIH). SARS-CoV-2 strain WA1 (NR-52281; BEI Resources) was propagated in Vero cells unless specified otherwise [6]. For viral stocks, cells were infected at a multiplicity of infection (MOI) of 0.01 and cultured for 48 h. At that point, cells were harvested, homogenized, subjected to a single freeze-thaw cycle, and then combined with the culture supernatant followed by centrifugation (3000 rpm, 10 min). The viral titers of the final supernatant (after serial dilution) was determined by plaque forming assay. All work with SARS-CoV-2 was performed under BSL3 conditions in a facility with negative pressure and PPE that included Tyvek suits and N95 masks for respiratory protection.

### 2.3. Viral irradiation with solar simulated UV light (sUV)

A KW large area light source solar simulator, model 91293, from Oriel Corp. (Stratford, CT) was used, equipped with a 1000W xenon arc lamp power supply, model 68920, and a VIS-IR band pass blocking filter plus either an atmospheric attenuation filter (output 290–400 nm plus residual 650–800 nm for solar simulated light) [19,20]. For viral irradiation, viral stocks were diluted >1:100 in PBS and irradiated in a sealed UV-transparent cuvette [BrandTech^™^ BRAND^™^ UV-Cuvets, providing transparency from 230 to 900 nm, widely used for DNA, RNA and protein analysis (BrandTech^™^ 759170, Fisher Scientific)]. The cuvette was inserted into a fully UV-transparent scintillation counter vial (Wheaton ‘180’ low-potassium glass, SigmaAldrich Z253081). The UV output was quantified using a dosimeter from International Light Inc. (Newburyport, MA), model IL1700, with an SED240 detector for UVB (range 265–310 nm, peak at 285 nm) or a SED033 detector for UVA (range 315–390 nm peak 365 nm) at a distance of 365 mm from the source, which was used for all experiments. In order to avoid artifactual thermal effects of photon exposure on viral activity, cuvettes were placed on ice during irradiation. At 365 mm from the source, total solar UV intensity was 5.34 mJ/cm^2^ s (UVA) and 0.28 mJ/cm^2^ s (UVB).

### 2.4. HCoV-NL63 plaque forming assay and viral RNA quantification

A published standard procedure was followed [15,21]. For HCoV-NL63, target cells (CaCo-2 or Calu-3) were seeded in 6-well plates at approximately 4 × 10^5^ cells per well and incubated until the monolayer was 80–90% confluent. Prior to infection, cells were washed with phosphate buffered saline (PBS). Virus inoculum (MOI=0.01) in 500 μL of growth media supplemented with 2% horse serum (with standard penicillin/streptomycin and L-glutamine supplementation) was added to each well. Viral entry was performed by incubation at 4°C for 30-60 min with gentle agitation followed by 1 h incubation in 33°C, 5% CO_2_. Then, inoculum was removed and cells were washed twice with PBS and replaced by 2 mL of normal growing media. After infection, cells were washed twice with PBS and placed in the incubator and cultured in normal growth media. Once plaques appeared (∼5-7 d post infection), cells were fixed with 10% neutral buffered formalin for 30 min at room temperature and stained with 1% crystal violet in 20% methanol for 20 min. Then, cells were washed several times with water, and plaques were counted and representative pictures taken at 10x magnification using an inverted microscope (Nikon Instruments, Melville, NY). In addition, viral RNA was extracted from cells and the respective culture supernatant with the QIAamp Viral RNA Mini Kit (QIAGEN, Hilden, Germany). One step RT-qPCR for HCoV-NL63 with absolute virus RNA quantification was performed using the following primer/probe set as published before [22]:

forward primer – 5′-ACGTACTTCTATTATGAAGCATGATATTAA-3′

reverse primer – 5′-AGCAGATCTAATGTTATACTTAAAACTACG-3′

probe – FAM-5′-ATTGCCAAGGCTCCTAAACGTACAGGTGTT-3′-NFQ-MGB

Briefly, RT-qPCR was carried out in a 20 µL reaction mixture with extracted RNA and One step RT-qPCR 2x Master Mix containing ROX as a passive reference dye (Gold Biotechnology, St. Louis, MO) and 300 nM forward and reverse primers and 200 nM MGB probe. Amplification and detection were performed in ABI 7500 system (Applied Biosystems, Foster city, CA) under the following conditions: first strand cDNA synthesis at 42°C for 30 min; initial denaturation/RT inactivation at 95°C for 3 min; denaturation at 95°C for 10 sec and annealing/extension at 55°C for 30 sec followed by 45 sec for data acquisition at 72°C. During amplification, the ABI PRISM 7500 sequence detector monitored real-time PCR amplification by quantitative analysis of the fluorescence emissions. The reporter dye (FAM) signal was measured against the internal reference dye (ROX) signal to normalize the signals for non-PCR-related fluorescence fluctuations that occur from well to well. The cycle threshold (Ct) represented the refraction cycle number at which a positive amplification was measured and was set at ten times the standard deviation of the mean baseline emission calculated for PCR cycles 3 to 15. Genomic RNA from HCoV-NL63 was used as a positive control.

### 2.5. SARS-CoV-2 plaque forming assay and viral RNA quantification

The quantification of infectious SARS-CoV-2 has been published before [18]. Target cells (Vero or Calu-3) were infected in triplicates at an MOI of 0.005 (high titer) or 0.001 (low titer). Briefly, cells were incubated with SARS-CoV-2 for 2 h and subsequently overlaid with 1% methylcellulose in culture medium. After 3-4 days, the cells were fixed in 10% neutral buffered formalin for 30 min, washed under tap water, and stained with 1% crystal violet. The number of plaques was counted on a light table. Alternatively, infection of cells was determined by measuring the amount of viral RNA. Cells were lysed in Trizol followed by RNA extraction with the RNAeasy kit (Qiagen). After reverse transcription, cDNA corresponding to the gene encoding the SARS-CoV-2 spike protein was quantified by qPCR with the Perfecta FastMix (QuantaBio) using:

forward primer (SARS-CoV-2) 5′-GCTGGTGCTGCAGCTTATTA-3′

reverse primer (SARS-CoV-2) 5′-AGGGTCAAGTGCACAGTCTA-3′

at an annealing temperature of 60 ºC. For normalization, *GAPDH* expression was measured using the following primers:

forward primer (*GAPDH*) 5′-TGGTGAAGGTCGGTGTGAAC-3′

reverse primer (*GAPDH*) 5′-CCATGTAGTTGAGGTCAATGAAGG-3′.

### 2.6. Human Stress & Toxicity PathwayFinder RT^2^ Profiler™ gene expression array analysis of infected host cells

Seven days post infection of Calu-3 host cells with either HCoV-NL63 (MOI=0.01) or HCoV-NL63 exposed to sUV (UVB portion: 706 mJ/cm^2^), total mRNA from host cells was isolated using the RNeasy Mini kit (Qiagen, Valencia, CA) following our published standard procedures. Reverse transcription was then performed using the RT^2^ First Strand kit (Qiagen) from 500 ng total RNA. For gene expression array analysis, the human Stress & Toxicity PathwayFinder RT^2^ Profiler™ technology (Qiagen), assessing expression of 84 stress response-related genes, was used as published before [23,24]. Quantitative PCR was run using the following conditions: 95 °C (10 min), followed by 40 cycles at 95 °C (15 s) alternating with 60 °C (1 min) (Applied Biosystems, Carlsbad, CA). Gene-specific products were normalized to a group of 5 housekeeping genes (*ACTB, B2M, GAPDH, HPRT1, RPLP0)* and quantified using the comparative ΔΔCt method (ABI PRISM 7500 sequence detection system user guide). Expression values were averaged across at least three independent array experiments, and standard deviation was calculated for graphing and statistical analysis as published before.

### 2.7. Individual RT-qPCR analysis

Total cellular mRNA was isolated using the Qiagen RNeasy Mini Kit (Qiagen, Gaithersburg, MD) according to the manufacturer’s protocol as published by us before [24]. Human primer probes [*CCL3* (Hs_00234142_m1), *CSF2* (Hs_00929873_m1), *HSPA6* (Hs_00275682_s1), *IL1B* (Hs_00174097_m1), *IL6* (Hs_00985639_m1), *SOD2* (Hs_00167309_m1), *TNF* (Hs_00174128_s1), and *RSP18 (*housekeeping gene; Hs_01375212_g1)], were obtained from ThermoFisher Scientific (Waltham, MA). After cDNA synthesis, quantitative PCR reactions were performed as follows: 10 min (95 °C) followed by 15 sec (95 °C), 1 min (60 °C), 40 cycles, using the ABI7500 Real-Time PCR System (Applied Biosystems, Foster City, CA). Amplification plots were generated, and Ct values were recorded as published before [24].

### 2.8. Statistical analysis

Unless stated differently, data sets were analyzed employing analysis of variance (ANOVA) with Tukey’s posthoc test using the GraphPad Prism 9.1.0 software (Prism Software Corp., Irvine, CA); in respective bar graphs (analyzing more than two groups), means without a common letter differ (p < 0.05) as published before [24]. For bar graphs comparing two groups only, statistical significance was calculated employing the Student’s two-tailed t-test, utilizing Excel (Microsoft, Redmond, WA). Experiments were performed in sets of at least three independent repeats. The level of statistical significance was marked as follows: *p < 0.05; **p < 0.01; ***p < 0.001.

## 3. Results

### 3.1. Solar simulated UV exposure of HCoV-NL63 blocks subsequent viral infection and replication in Calu-3 human epithelial lung cells

First, we examined the feasibility of UV-inactivation of a pathologically relevant coronavirus by employing a single dose of solar simulated UV light using a commercial xenon light source with quantified spectral power distribution (Fig. 1A). To this end, we exposed human coronoavirus NL63 (HCoV-NL63) in PBS to a high dose of sUV [equivalent to approximately 6 minimal erythemal doses (MEDs; UVA: 13.46 J/cm^2^; UVB: 706 mJ/cm^2^)] and subsequently used it to infect Calu-3 target cells for 7 days [2, 6, 8]. We used unexposed virus as controls. Strikingly, sUV pre-exposure strongly suppressed viral infectivity of target cells as demonstrated by quantitative plaque assay analysis, indicating that sUV exposure caused a more than 8-fold decrease in viral infectivity (Fig. 1B).

**Figure 1.**
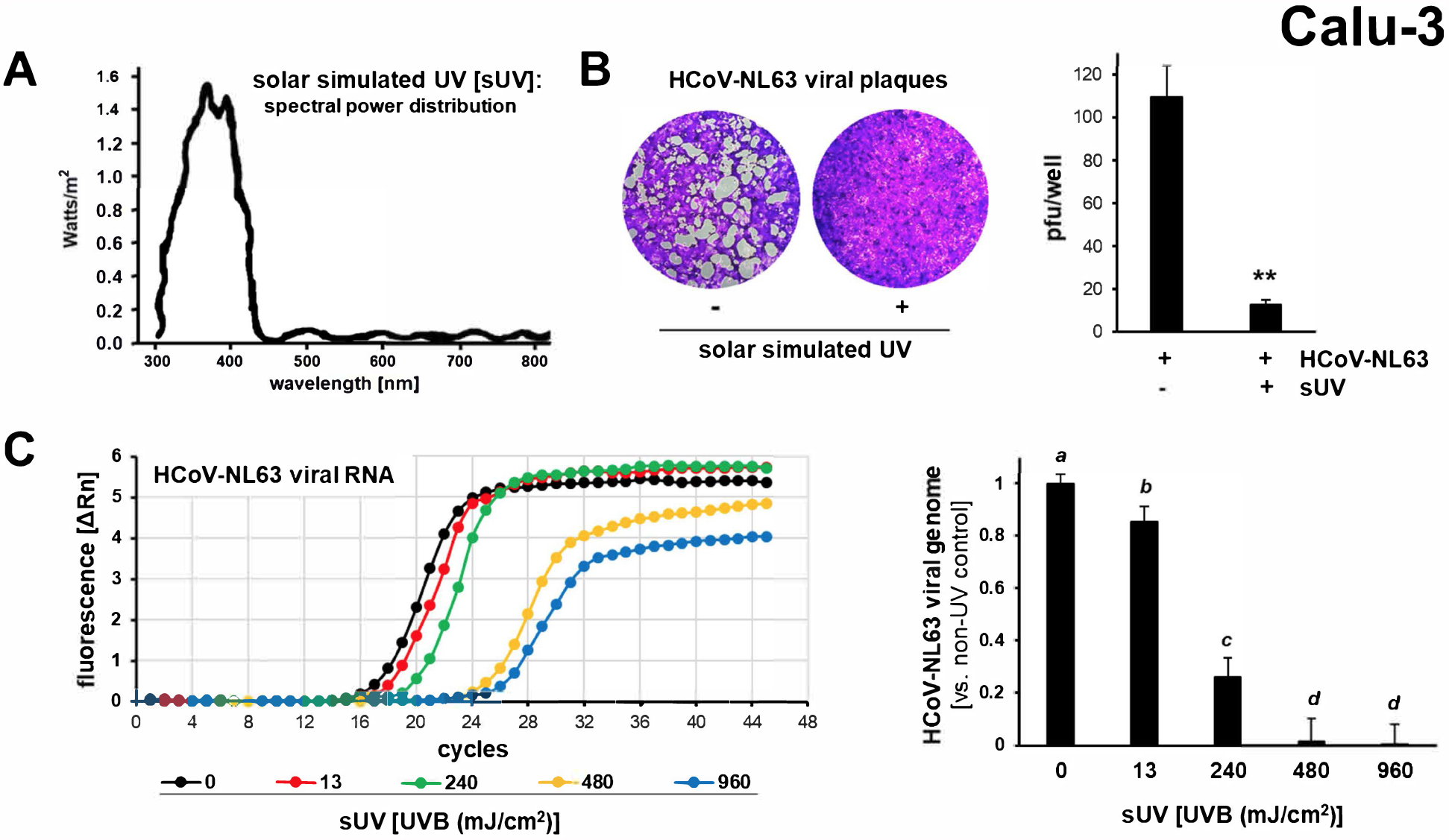
Solar simulated UV pre-exposure antagonizes HCoV-NL63 viral infectivity targeting Calu-3 human epithelial lung cells. Virus in PBS was exposed to sUV or left unexposed followed by Calu-3 target cell infection (0.01 MOI) and post infection culture over 7 days followed by analysis. (A) Spectral power distribution (irradiance) of the solar simulator light source equipped with appropriate cut-off filter (sUV: UVB + UVA, solid black line). (B) Plaque assay after viral exposure to sUV (UVB portion: 706 mJ/cm^2^) as visualized by light microscopy (10 x magnification); bar graph summarizes numerical data. (C) RT-qPCR of viral genome replication in target cells [left panel: amplification curves as a function of sUV dose (UVB portion as indicated); right panel: bar graph summarizing numerical data].

Next, we examined the dose-response relationship characterizing the inhibition of HCoV-NL63 viral replication (induced by sUV pre-exposure) by one step RT-qPCR analysis of the genomic RNA copy number. We detected a significant inhibition at low sUV doses [UVA: 0.25 J/cm^2^; 13 mJ/cm^2^ UVB]. Viral inactivation of more than 98 % occurred at doses equal and above 480 mJ/cm^2^ UVB (UVA: 9.04 J/cm^2^; Fig. 1C).

### 3.2. Solar simulated UV exposure of HCoV-NL63 blocks subsequent infection of Caco-2 human epithelial colorectal cells

In order to explore sUV effects on HCoV-NL63 infectivity in another human target cell, we exposed the virus (in PBS) to a high dose of sUV [equivalent to approximately 6 MEDs (UVA: 13.46 J/cm^2^; UVB: 706 mJ/cm^2^)] and subsequently infected Caco-2 epithelial colon cells (Fig. 2). As observed before with Calu-3 cells (Fig. 1), our quantitative plaque assay analysis showed that the suppression of viral infectivity of Caco-2 target cells by sUV exposure caused a more than 4-fold decrease in plaque formation (Fig. 2A). Likewise, our dose response analysis by RT-qPCR of genomic RNA copy numbers indicated that sUV exposure caused a pronounced suppression of HCoV-NL63 viral replication at doses as low as 240 mJ/cm^2^ UVB (UVA: 4.52 J/cm^2^; Fig. 2B).

**Figure 2.**
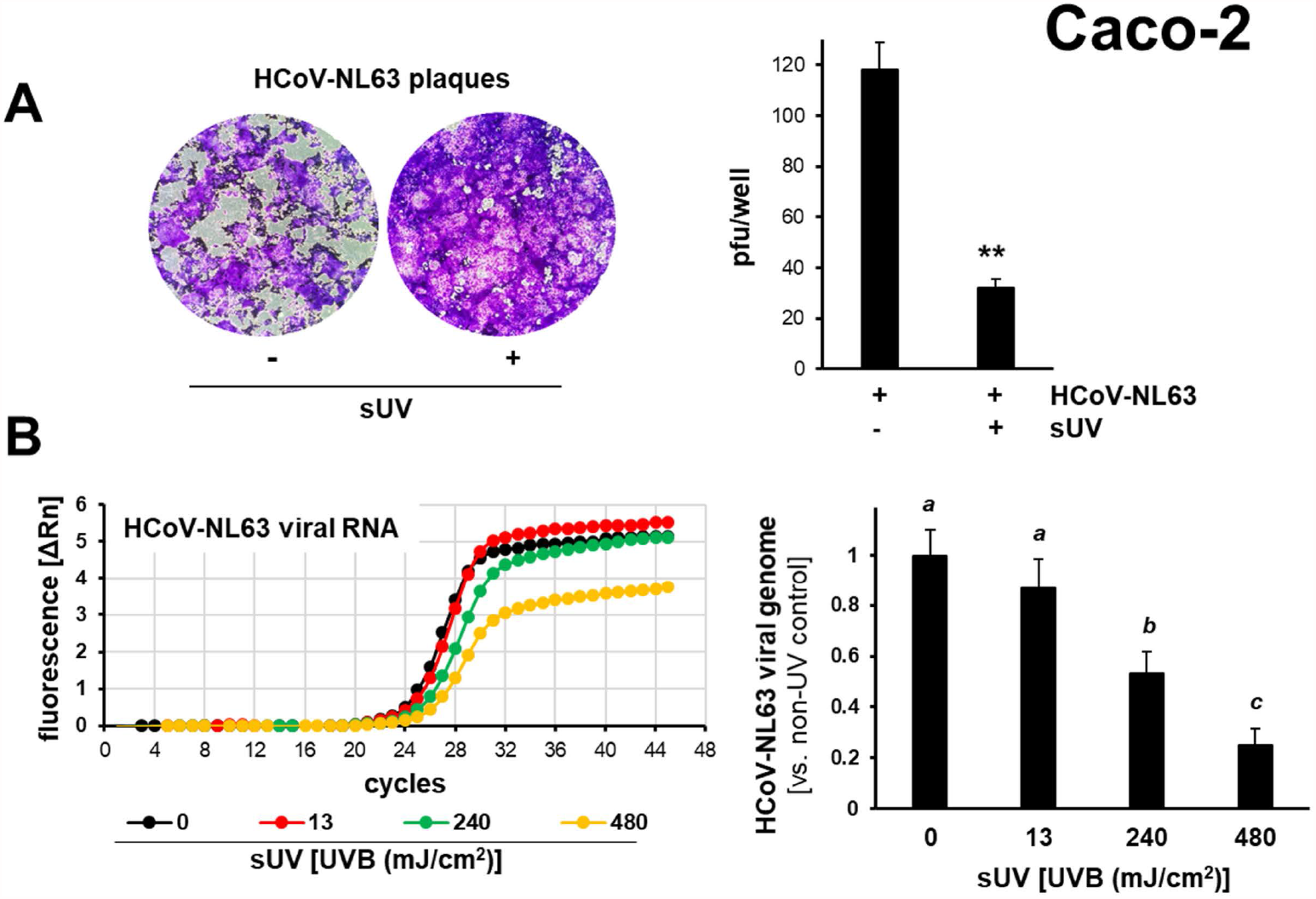
Solar simulated UV pre-exposure antagonizes HCoV-NL63 viral infectivity targeting Caco-2 human epithelial colorectal cells. Virus in PBS was exposed to sUV or left unexposed followed by Caco-2 target cell infection (0.01 MOI) and post infection culture (7 days) followed by analysis. (A) Plaque assay after viral exposure to sUV (UVB portion: 706 mJ/cm^2^) as visualized by light microscopy (10 x magnification); bar graph summarizes numerical data. (B) RT-qPCR detection of viral genome replication in target cells; left panel: amplification curves (as a function of sUV-dose); right panel: bar graph summarized numerical data.

### 3.3. Stress response gene expression array analysis confirms solar UV-induced inhibition of HCoV-NL63 infectivity targeting Calu-3 human epithelial lung cells

Next, the cellular stress response of Calu-3 human epithelial lung cells, elicited by infection with either mock-irradiated or sUV pre-exposed HCoV-NL63, was examined at the gene expression level using the RT^2^ Human Stress and Toxicity PathwayFinder™ PCR Array technology. To this end, we infected Calu-3 target cells with sUV or mock-treated virus (doses as in Figs. 1, 2) and profiled the gene expression at the end of the experiment. We observed global HCoV-NL63-induced expression changes (antagonized by viral pre-exposure to sUV) as depicted by Volcano plot (Fig. 3). As expected, HCoV-NL63 viral infection caused a pronounced upregulation of stress response gene expression including genes encoding key regulators of inflammatory signaling (such as *CSF2, TNF, IL1B, IL1A, CCL3, CXCL10, NFKBIA*, and *IL6*), oxidative stress defense (such as *SOD2*), and heat shock response (such as *HSPA6*; Fig. 3). In contrast, after viral sUV-exposure performed pre-infection, most of these infection-associated expression changes were either attenuated or completely obliterated, an observation consistent with pronounced suppression of HCoV-NL63 viral infectivity as a consequence of sUV-exposure. Likewise, HCoV-NL63 viral infection-induced expression changes causing downregulation of specific apoptotic modulators including *BCL2L1, EGR1, CASP8*, and *CASP1*, proliferation markers such as *PCNA*, and heat shock response factors such as *HSPA4, HSPH1*, and *HSP90AA2P* were completely absent in samples obtained from cells exposed to the pre-irradiated virus. Strikingly, expression of seven specific genes (*CDKN1A, CYP1A1, MDM2, HMOX1, RAD50, HSPA1L*, and *E2F1*) was modulated uniquely in response to exposure to sUV-preirradiated HCoV-NL63, a finding consistent with gene expression changes responsive to sUV-induced chemical damage to viral components (including ribonucleic acids, proteins, and lipids) [1–3].

**Figure 3.**
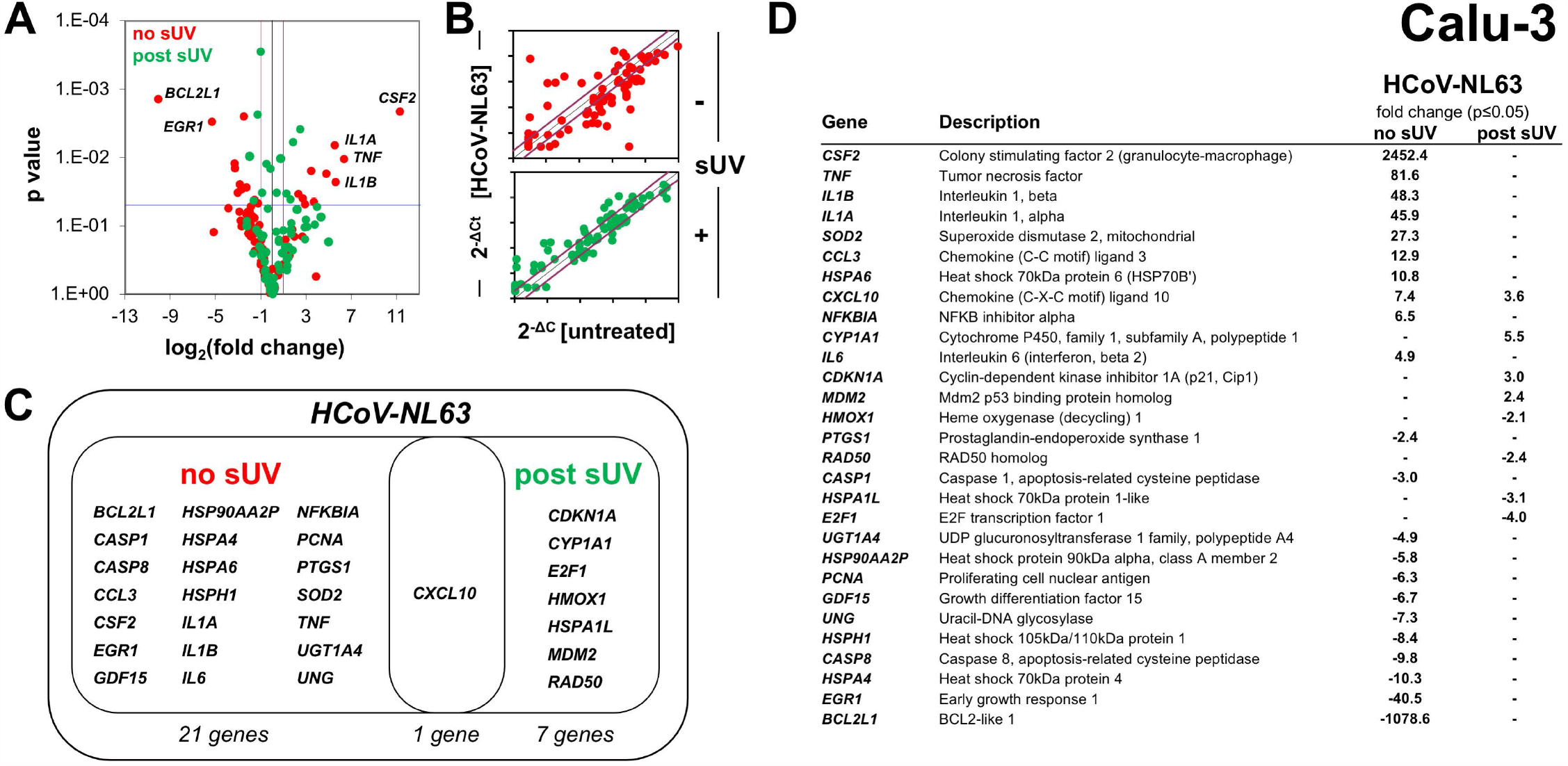
Solar simulated UV pre-exposure of HCoV-NL63 prevents stress response gene expression elicited in Calu-3 human epithelial lung target cells. Treatments were performed as detailed in Fig. 1. (A) Target cell stress response [control (HCoV-NL63) versus sUV (UVB portion: 706 mJ/cm^2^) pre-exposed virus] assessed by *RT*^*2*^ *Profiler*^*TM*^ *Stress and Toxicity Pathway* gene expression array analysis [volcano plot depiction: p value over log2 (fold expression change)]. (B) Scatter plot depiction comparing expression changes elicited by untreated control virus (top panel) or sUV pre-exposed virus (bottom panel). (C) Venn diagram depicting expression changes induced by mock-irradiated virus (control) versus sUV pre-irradiated virus. (D) Tabular summary of numerical values specifying gene expression changes at the mRNA level (p<0.05).

### 3.4. Dose-response relationship of solar simulated UV-induced inhibition of SARS-CoV-2 infectivity targeting Vero and Calu-3 mammalian cells

After demonstrating HCoV-NL63 coronavirus inactivation by sUV at an environmentally relevant dose level, we examined whether sUV-inactivation might also be applicable to SARS-CoV-2. To this end, we exposed the virus with a dose range of sUV, subsequently infected Vero monkey epithelial cells at two different multiplicities of infection (MOIs, high versus low titer), and measured the number of infectious virions three days later by plaque forming assay. Strikingly, as observed with HCoV-NL63, sUV exposure caused a pronounced suppression of viral infectivity. This antiviral effect, observable over a broad range of sUV doses, followed an exponential decay curve with an effective ED_50_ (sUV dose diminishing SARS-CoV-2 viral infectivity by 50%) approximating 55 mJ/cm^2^ (low titer) and 62 mJ/cm^2^ (high titer) (Fig. 4A).

**Figure 4.**
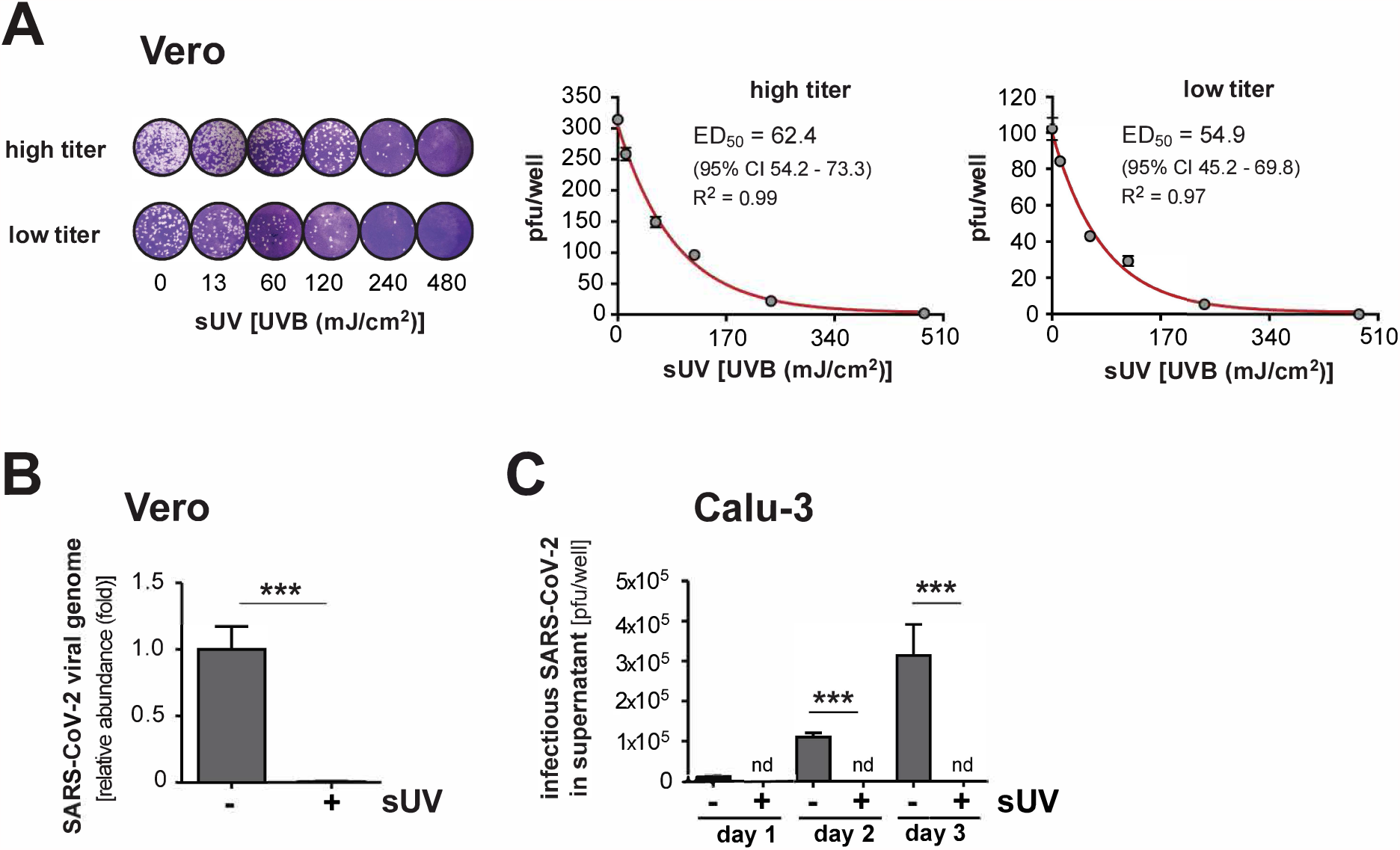
Solar simulated UV exposure of SARS-CoV-2 antagonizes subsequent viral infection and replication in African green monkey Vero and Calu-3 human epithelial lung cells. (A) SARS-CoV-2 was sUV-irradiated (UVB portion: up to 480 mJ/cm^2^; or remained unirradiated) in PBS and subsequently used to infect Vero cells at two different MOIs (high versus low titer). Dose response of plaque formation as a function of sUV pre-exposure dose was assessed; a representative experiment (left panel, top and bottom rows) and quantification (right panels) are depicted. (B) Detection of viral genome replication in Vero cells with quantification of viral RNA after infection using mock or sUV pre-irradiated virus (UVB portion: 1010 mJ/cm^2^) as assessed by RT-qPCR after 24 h. (C) Infection of Calu-3 cells with SARS-CoV-2 [sUV pre-exposed (UVB portion: 706 mJ/cm^2^) versus unirradiated virus]. The presence of infectious virions in the supernatants was quantified over the course of three days post infection by RT-qPCR (nd: not detectable).

Next, we tested feasibility of achieving complete inhibition of SARS-CoV-2 replication by high dose sUV [UVB portion: 1010 mJ/cm^2^, a maximum dose level similar to the one used in the HCoV-NL63-directed dose-response experiments (Fig. 1C)]. To this end, we pre-exposed SARS-CoV-2 to sUV and measured the amount of viral RNA (corresponding to the region of the viral genome encoding the S protein) by RT-qPCR analysis. Indeed, complete inhibition was achieved at that dose (Fig. 4B). We obtained similar results for sUV-exposed SARS-CoV-2 infections of Calu-3 human lung epithelial target cells with viral load in supernatants being monitored over three days by RT-qPCR (Fig. 4C). Taken together, we conclude that SARS-CoV-2 is sensitive to sUV suggesting viral inactivation at environmentally relevant exposure levels.

### 3.5. Solar simulated UV exposure of SARS-CoV-2 prevents stress response gene expression elicited by viral infection of Calu-3 human epithelial lung cells as detected by array analysis

Next, to determine Calu-3 human epithelial lung cell stress response gene expression elicited by SARS-CoV-2 as a function of viral pre-exposure to sUV, we employed expression analysis using the Human Stress and Toxicity PathwayFinder™ PCR Array technology. To this end, we infected Calu-3 target cells with sUV or mock-treated virus as outlined before, followed by comparative gene expression profiling at the end of the experiment. We observed multiple SARS-CoV-2-induced expression changes (antagonized by viral pre-exposure to sUV) as shown in the Volcano plot depiction [displaying statistical significance (P value) versus magnitude of change (fold change)] (Fig. 5). SARS-CoV-2 infection caused a pronounced upregulation of stress response gene expression including genes encoding key regulators of inflammatory signaling including *IL1A, IL1B, IL6, TNF, CCL3, CXCL10, CSF2, and NFKBIA*, oxidative stress defense such as *SOD2*, and heat shock response such as *HSPA6* (Fig. 5). In contrast, after infection with sUV-exposed virus, most of these infection-associated expression changes were either attenuated or completely obliterated, an observation consistent with pronounced suppression of SARS-CoV-2 infectivity as a consequence of sUV-exposure. Remarkably, these expression changes closely mirrored those observed in response to HCoV-NL63 infection that occurred with or without viral exposure to sUV (Fig. 3).

**Figure 5.**
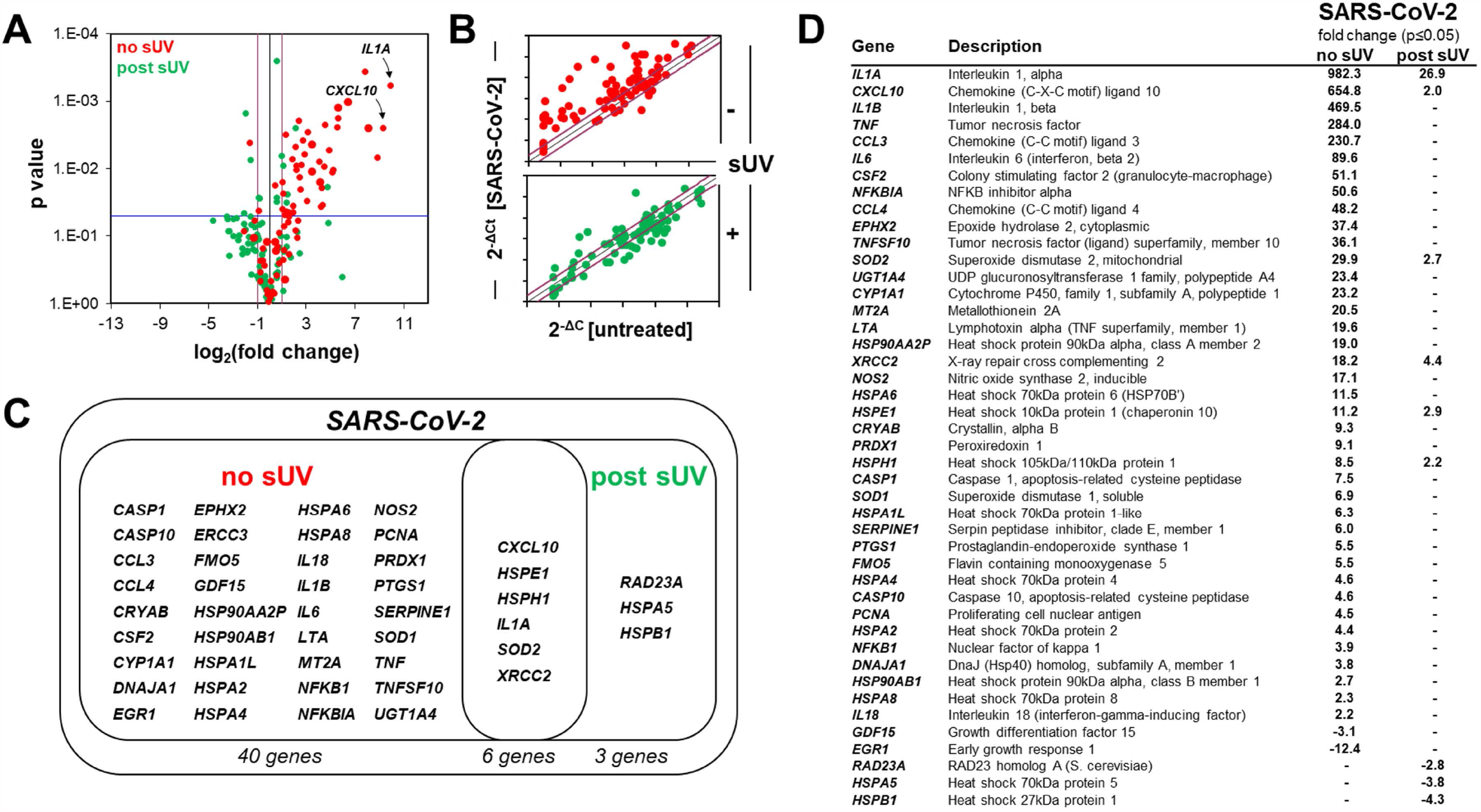
Solar simulated UV pre-exposure of SARS-CoV-2 prevents stress response gene expression elicited in Calu-3 human epithelial lung target cells. Treatment and analysis were performed as detailed in Fig. 3. (A) Target cell stress response [control (SARS-CoV-2) versus sUV (UVB portion: 706 mJ/cm^2^)-preirradiated virus] assessed by *RT*^*2*^ *Profiler*^*TM*^ *Stress and Toxicity Pathway* gene expression array analysis [volcano plot depiction: p value over log2 (fold expression change)]. (B) Scatter plot depiction comparing expression changes elicited by untreated control virus (top panel) or sUV pre-exposed virus (bottom panel). (C) Venn diagram depicting expression changes induced by mock-irradiated virus (control) versus sUV pre-irradiated virus. (D) Tabular summary of numerical values of gene expression changes at the mRNA level (p<0.05).

Likewise, we observed a striking similarity between the gene expression changes elicited by HCoV-NL63 and SARS-CoV-2 (and blocked by viral sUV pre-exposure), modulating redox, inflammatory, and proteotoxic stress responses in Calu-3 human epithelial lung cells (Fig. 6). Specifically, sUV-induced (UVB portion: 706 mJ/cm^2^) viral inactivation was apparent from independent RT-qPCR assessment of mRNA levels (‘no sUV’ versus ‘sUV’) interrogating genes encoding key regulators of redox (*SOD2*), inflammatory (*IL1B, TNF, CCL3, IL6, CSF2*), and proteotoxic (‘heat shock’; *HSPA6*) stress responses in Calu-3 target cells as detailed above. Thus, our data suggest that similar to HCoV-NL63, sUV exposure of SARS-CoV-2 interrupts the viral life cycle causing suppression of viral replication and virus-induced inflammatory and cellular stress responses in mammalian target cells.

**Figure 6.**
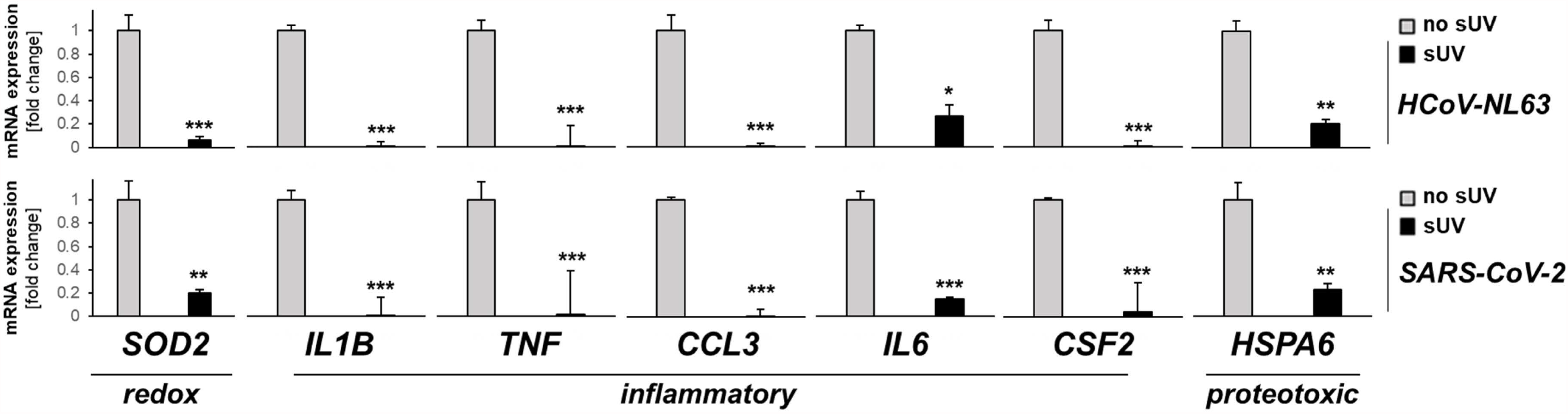
Comparative analysis of redox, inflammatory, and proteotoxic stress response gene expression in Calu-3 human epithelial lung cells elicited by HCoV-NL63 and SARS-CoV-2 (with and without viral sUV pre-exposure). Gene expression as assessed by single RT-qPCR quantification in virus-exposed target cells as a function of viral pre-exposure [‘no sUV’ versus ‘sUV’ (UVB portion: 706 mJ/cm^2^)]. Bar graphs depict fold change (‘sUV’ versus ‘no sUV’) normalized to housekeeping gene expression (*RPS18*; gray bar: no sUV pretreatment; black bar: sUV-pretreatment).

## 4. Discussion

Identification and mechanistic exploration of environmental factors that might determine coronavirus infectivity are of significant interest with relevance to both basic molecular research and public health-related preventive and interventional investigations [2]. Here, we have explored for the first time the effects of full spectrum (UVA + UVB) solar ultraviolet radiation on coronavirus infectivity and demonstrate that sUV inactivates HCoV-NL63 and SARS-CoV-2 coronaviruses at environmentally relevant doses. First, we observed that exposure of HCoV-NL63 and SARS-CoV-2 to sUV (performed at acute dose levels relevant to human populations worldwide) blocks subsequent viral infection and replication in relevant primate target cells [human: Calu-3 lung epithelial, Caco-2 colorectal epithelial; monkey: Vero kidney epithelial (Figs. 1, 2, 4)]. Blockade of viral infectivity in response to sUV pre-exposure was also confirmed using stress response gene expression profiling in array (Figs. 3, 5) and independent RT-qPCR format (Fig. 6) elicited in Calu-3 target cells by coronavirus infection (HCoV-NL63 and SARS-CoV-2).

Remarkably, dose levels used throughout this pilot study are representative of terrestrial ground level exposure suggesting environmental relevance, and significant coronavirus inactivation was detectable even at low exposure levels expected to be beneath the cutaneous sunburn-inducing threshold (Figs. 1, 2, 4) [2,6,8]. In this context, it is remarkable that recent research has already indicated that ground level solar UV displays significant virucidal effects targeting coronaviruses including SARS-CoV-2 [2,6,11,13]. However, the complexity of human exposure levels to solar UV as a function of solar zenith angle, seasonality, spectral distribution, and latitude remain to be addressed before any firm conclusions relevant to human populations can be drawn. Specifically, the anti-viral activity of specific spectral components of sUV remains to be determined since the light source employed in our prototype studies emitted full spectrum simulated solar UV, and the action spectrum of virus inactivation by solar UV remains largely undefined. For example, it is possible that the UVA portion of ground level sUV significantly contributes to the coronavirus-directed effects described by us [7]. It therefore remains to be seen if indirect impairment of viral structure and infectivity occurs by alternative mechanisms, such as UVA-driven photosensitization and oxidative stress (mediated by formation of reactive oxygen species including singlet oxygen), that might be operative in addition to direct inactivation of viral genomic RNA through nucleic acid base photodamage. It will also be interesting to explore potential mechanistic synergisms underlying virucidal effects that occur upon combined UVA and UVB as compared to separate spectral exposure. Likewise, experimental conditions used throughout our studies (including viral irradiation in PBS and exposure performed in cell culture medium) might limit the applicability of our conclusions in the context of relevant coronavirus transmission situations that involve more complex determinants of infectivity including the role air-borne and aerosol transmission and intermediate surface retention [6].

Addressing urgency and global scale of the unfolding SARS-CoV-2 pandemic requires an improved understanding of environmental factors that modify viral infectivity [2,6,8]. Taken together, our data suggest feasibility of sUV-induced viral inactivation targeting HCoV-NL63 and SARS-CoV-2 coronaviruses, a finding to be substantiated by future mechanistic exploration performed in more relevant *in vivo* exposure models.

## Abbreviations

MOI: multiplicity of infection
sUV: solar simulated ultraviolet light
UV: ultraviolet.

## Acknowledgement

The authors like to acknowledge technical support provided by Jennifer L. Uhrlaub.

